# GEMspa: a Napari plugin for analysis of single particle tracking data

**DOI:** 10.1101/2023.06.26.546612

**Authors:** Sarah Keegan, David Fenyö, Liam J. Holt

## Abstract

The cellular environment is crowded with macromolecules and far from thermodynamic equilibrium. This active, crowded environment influences biochemical reactions and the formation of cellular structures such as membraneless organelles. These physical properties can change during normal physiology and in disease states such as neurodegenerative diseases and cancer, impacting cell behavior and function. Therefore, it is crucial to develop methods to characterize these properties. Microrheology is the inference of physical properties from the motion of tracer particles embedded within a material. This technique requires single particle tracking (SPT) and analysis of these tracks through the framework of soft-condensed matter physics. Analysis of SPT data can be challenging due to the lack of comprehensive user-friendly software tools. To address this, we introduce GEMspa, a software package implemented as a plugin for the open source image analysis platform, napari. GEMspa provides a GUI for a commonly used localization/tracking algorithm (via Trackpy), and a suite of methods to extract basic parameters describing particle motion. This platform aims to streamline the workflow of data analysis steps and allow researchers to visualize and optimize parameters for high-quality results, thereby making microrheology accessible to non-expert scientists.

## INTRODUCTION

The cell interior is crowded with macromolecules that occupy approximately 15-30% of its total volume^1, 2^. This characteristic is not often accounted for in dilute *in vitro* solutions^3^, however biological molecules have evolved to function in an environment “concentrated” with macromolecules. Macromolecules that span the mesoscale size range (diameters from ∼10 - 1000 nm) are responsible for most of the excluded volume within the cell, that is, they physically exclude other molecules from the volume that they occupy. This exclusion is size-dependent; smaller molecules can move more freely since they can “fit between’’ larger molecules much as a motorcyclist can weave in and around traffic, while molecules of similar size or larger than the crowder are restricted in their movement^1, 4^. Molecules can experience a “traffic jam” when crowding is exacerbated. For example, inducing a sudden reduction in cellular volume by severe osmotic stress leads to reduced biochemical reaction rates and stalling of cellular processes^5^. While crowding can hinder reactions as a result of reduced diffusivity, it can also promote reactions that result in depleting the total excluded volume of a system through the depletion attraction effect^5, 6^. The association of molecules, which typically results in a reduction of entropy, can actually increase the entropy of a system as a whole since it makes more space available to the crowder. An example of this effect can be found in the formation of liquid-liquid phase separated (LLPS) condensates, which can be regulated through changes in crowding both *in vitro* and *in vivo*^7^. Recently, it was found that the phase separation of WNK kinases in response to hyperosmotic crowding is required for their regulation of cell volume increase^8^. In fact, an array of multivalent proteins reversibly phase-separate upon perturbation of crowding^9^. Membraneless compartments within the cell, such as nucleoli, Cajal bodies, nuclear speckles, stress granules and p-bodies all display properties of liquid-like condensates and may therefore be regulated by changes in crowding.

The biophysical properties of the cell can be altered in stress and disease states. For example, the formation of protein aggregates is a hallmark of many neurodegenerative diseases. They have been found to undergo maturation, a transition from a dynamic state of reversible liquid-liquid phase separation to an irreversible state where they form fibrillar aggregates^10^. Modeling of molecular condensation associated with increased crowding over time resulted in a slow build up and rapid onset of protein aggregation^11^. While the formation of these aggregates is still not well understood, it is possible that changes in the biophysical properties of the cell plays a role. In contrast, decreased crowding can also be a problem. Arresting the cell cycle and allowing yeast cells to become oversized results in cytoplasmic dilution due to their inability to scale the synthesis of proteins and RNA^12^. Yeast cells undergoing replicative aging as well as senescing human cells both suffer from increased cell size. Dilution alters biophysical properties of the cytoplasm and affects chemical reaction rates and molecular assembly. Exploring how these properties are associated with senescence and aging is an important avenue of investigation.

Cancer is a disease characterized by a disruption in the surrounding tissue: solid stresses can become high enough to compress blood and lymphatic vessels and interstitial fluid pressure and tissue stiffness increase. These physical traits activate mechanosensitive signaling pathways that contribute to tumor malignancy^13^. Growth-induced mechanical pressure has been found to reduce protein expression as a result of decreased molecular diffusion^14^. How cancer cells adapt and proliferate in compressive environments that can alter their biophysical properties remains an open question for investigation.

Techniques from microrheology can be used to study cellular biophysical properties. It is the study of the flow and deformation of matter on a microscopic scale. Cells exhibit rheological properties of both solids (elasticity) and fluids (viscosity) and are therefore said to be viscoelastic materials^15^. Microrheology methods measure the movement of tracer particles embedded within a medium of interest to extract its properties^16^. Active microrheology methods, such as optical or magnetic tweezers and atomic force microscopy perturb a tracer particle and observe its response^15, 17, 18^. Passive microrheology methods observe a particle’s movement with no external perturbation. These include bulk methods such as dynamic light scattering^19^ and its extension, diffusing wave spectroscopy^20^, as well as single particle tracking (SPT)^21, 22^. In SPT, probe particles are imaged over time at a high frame rate and particle positions are localized and tracked with specialized algorithms. These tracks are then analyzed to understand particle motions and the properties of the substance in which they are embedded. Unlike bulk methods, this technique allows for investigation into the heterogeneity of a system through examining single particle traces.

The analysis steps for SPT consist of localization, linking and quantification of motion. Many algorithms for particle localization exist^22–25^. Generally, the main steps are to perform image pre-processing to maximize the signal to noise ratio, isolate bright spots in the image, and then locate particle positions within these bright spots with sub-diffraction accuracy. This is performed either by iteratively fitting a point spread function to the diffraction limited spot or finding the intensity centroid. More recently, deep learning has also been applied to the task of particle localization with published success, especially in the context of super resolution microscopy^26–28^. The next step is linking particle positions from each frame into traces over time. This is non-trivial especially when particle density is high. It can become impossible if the distance between individual particles is less than the distance a single particle will travel between frames. Numerous algorithms also exist for this step, from the simple approach of linking nearest neighbors to more complex algorithms that take multiple frames and/or tracks into account and use mathematical optimization or statistical models to calculate the most likely combination of particle trajectories given the joint data^23, 29–34^. Factors such as the expected type of motion, properties of the particle intensity distribution, and false detection and birth/death rates can be taken into account for these calculations. Once particle positions have been linked, the analysis of particle motion can be performed. This generally encompasses fitting an applicable motion model^35^ to the track data to extract quantifiable parameters. One of the most common analysis methods is to extract the *diffusion coefficient* (D) from the mean square displacement (MSD), which can be calculated from a particle’s trajectory. This relation was described by Einstein^36^ in 1905 when he derived a theoretical formulation of Brownian motion. The equation states that the diffusion of a sphere moving through a fluid is inversely proportional to the sphere’s radius and the viscosity of the fluid. The equation is applicable at low inertia (low Reynold’s number, i.e. low turbulence of the fluid) relative to viscosity and when solutes are relatively small compared to the sphere. In his theory, a normalized Gaussian distribution provides the solution to the diffusion equation, and its variance, the MSD, scales linearly with time. The MSD is defined as the average squared distance a particle travels in a given increment of time (time-lag), where the average is taken for all time increments over the lifetime of the particle. Specifically, this formulation of the MSD is defined as the time-averaged MSD (TA-MSD), since ensemble averaging of the MSD (EA-MSD) can also be performed to extract the diffusion coefficient. The ensemble method averages over all distances traveled for all particles at a specific time. In the case of Brownian (random) motion, the MSD scales linearly with time-lag and the slope is proportional to the diffusion coefficient. A difference observed in the diffusion coefficient extracted from the TA or EA-MSD data would indicate a *non-ergodic* system, although statistical noise in the data should be taken into account in this comparison^22^. Ergodicity is the concept that at any time one can examine the data from an ensemble of particles and extract the same information as examining a single particle for a sufficiently long period of time. Ergodicity breaking indicates the presence of (non-random) anomalous motion and is a common feature found in SPT data from biological specimens^37–41^. The nature of anomalous motion can be found by applying a power law fit to the MSD and time-lag data to extract the *anomalous exponent* (α). The deviation of this exponent from one indicates motion that is sub-diffusive (less than 1) or super diffusive (greater than 1). A number of models exist for interpretation of anomalous motion found in SPT data. One such model is fractional Brownian motion (FBM), where there exists a correlation between successive increments of particle movement. This model can be applied in the context of anti-correlated motion resulting in sub-diffusion, which in living cells can be from the presence of macromolecular crowding and viscoelasticity^42, 43^.

Data analysis for SPT experiments can be challenging, both in choosing the correct algorithms and models and in finding easy to use and robust software tools. Often performing SPT data analysis will involve exporting data from a software application that implements localization and tracking to perform quantification of motion separately, complicating pipelines and separating the final step from the overall workflow. Quantification of motion also generally requires a researcher to hold higher than average computational skills, as tools with a graphical user interface (GUI) to backend code for applying models are lacking. We introduce GEMspa, a new software package for SPT data analysis. It is implemented as a plugin for the open source image analysis platform napari^44^. GEMspa provides a GUI for a python implementation^45^ of a commonly used and classical localization/tracking algorithm (Trackpy)^33^. It also provides a suite of methods to extract basic parameters describing particle motion. The aim of the software is to provide a unifying interactive platform for the real-time visualization of the results of localization, tracking and motion analysis, through the use of the layers feature of napari. GEMspa annotates points and tracks with informational properties for visualization of results so that parameter settings can be optimized. It also allows the user to perform all analysis steps within one application. GEMspa can be used for complete analysis from start to finish, giving a researcher an overall view of their data. It provides quality control measures as output to help the user evaluate results at each step, and its straightforward tab-oriented interface that follows the workflow of data analysis steps allows users to easily and quickly visualize and update parameters to obtain high quality results.

## RESULTS

### Overview

GEMspa is a napari plugin for analysis of data from single particle tracking microrheology experiments. To use GEMspa, first <u>install napari</u> and then GEMspa can be installed and accessed from the napari Plugins menu. The GEMspa GUI is designed to follow the workflow of analysis, allowing the researcher to set parameters and visualize results at each step. Once a video of particle movements is imaged, the workflow for SPT data analysis typically includes the following steps: (1) particle localization, (2) linking of localized particle positions into tracks, and (3) quantification of particle motion and inference of the underlying biophysical properties of the material under study (Figure 1a). Each step involves choices by the researcher on which algorithms or models to apply, and how to optimize parameter settings, as well as quality control of the output. The organization of the interface follows a tab-view design, allowing the researcher access to a distinct GUI for each step (Figure 1b, top right). The first tab, **New/Open** (Figure 1b, right panel), allows for import of imaging data in TIFF or ND2 file formats. It also gives an option to automatically create a napari labels layer to match the dimensions of the image (height/width), allowing the researcher to perform interactive masking to outline a region of interest. The native napari interface will by default create a labels layer to match the imported image, including the time dimension. Since this is not always the desired behavior, as only a single mask may be needed for the entire movie, we provide this convenience function. Finally, GEMspa allows for import of localization/tracking results from other software. This makes the quantification functionality of GEMspa available to researchers who have already performed localization and tracking separately. Currently, GEMspa supports import of results from three popular tracking algorithms: Mosaic^29^, Trackmate^46^ and Trackpy^45^. GEMspa provides a GUI interface for localization/tracking with Trackpy, but if a researcher prefers to implement this separately, for example as a stand-alone python script, Trackpy output can be imported here. The format for GEMspa tracking data is a simple tab-delimited text file with straightforward column names (Supplementary file 1). Any other outside localization and tracking results can also be imported to GEMspa by re-formatting the data to match the GEMspa file format.

**Figure 1.**
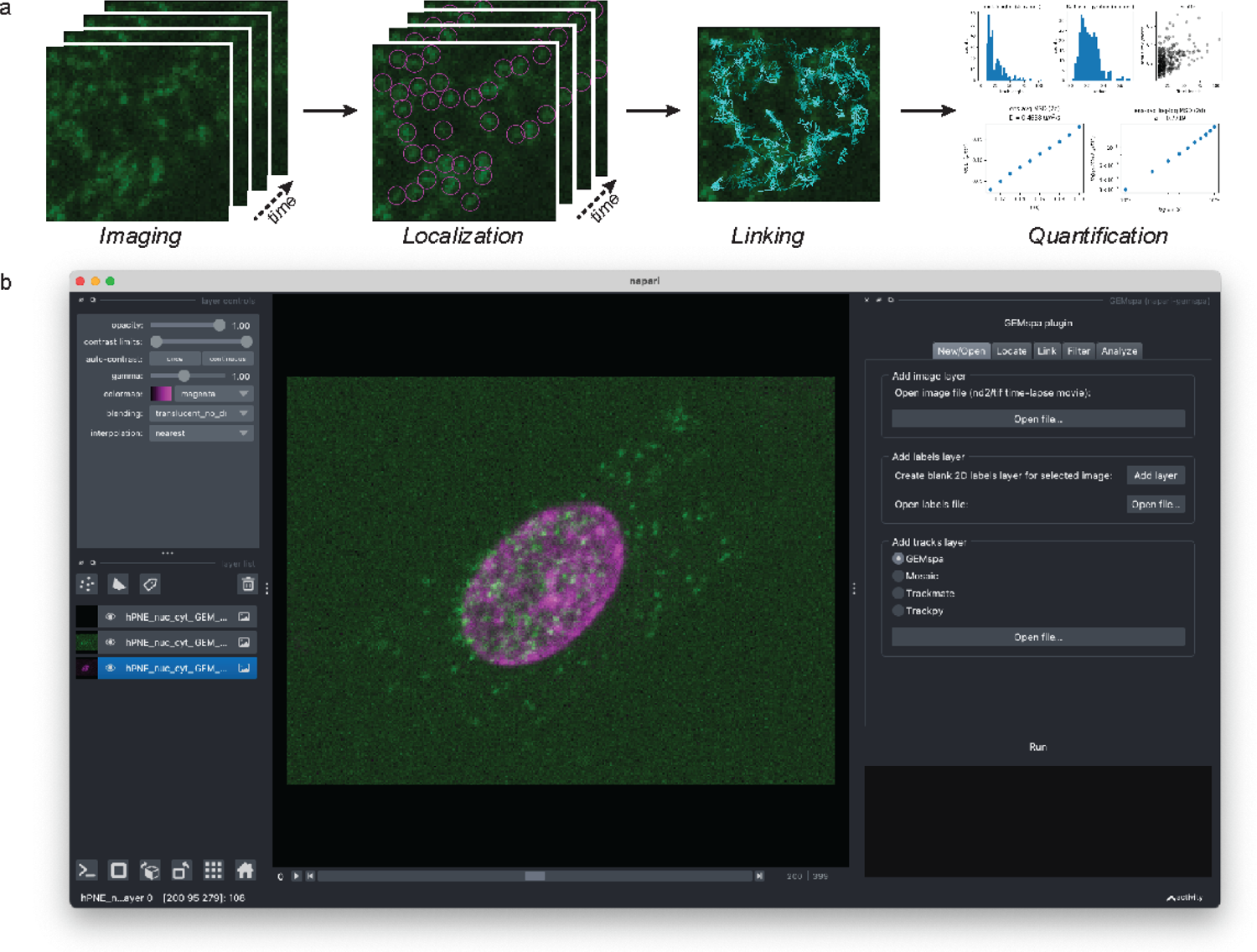
Overview of SPT Data Analysis and GEMspa plugin. (a) Cartoons illustrating the steps of SPT data analysis. (b) A screenshot of the GEMspa plugin contained within the napari application.

The subsequent tabs (Figure 1b, right panel) can be selected to navigate to GUI’s for performing localization, linking, filtering of linked tracks, and quantification of motion. Here, we will guide the reader through each step of analysis with an example movie of nucGEM particles^41^ diffusing in a hPNE (human pancreatic) cell (Figure 1b and Supplementary movie) imaged with a spinning-disk confocal at 100 Hz (100 frames per second) for 4 seconds. GEMs are genetically encoded multimeric nanoparticles based on naturally occurring homomultimeric scaffold proteins fused to fluorescent proteins^7^. GEMs use encapsulins as scaffolds, which assemble into bright, stable particles of a defined shape and size (20-40 nm diameter) and have been used as mesoscale probe particles by us and others^7, 12, 41, 47–49^. Specifically, nucGEMs were developed to probe the nucleoplasm and include a nuclear localization signal (NLS) which directs GEMs to assemble within the nucleus. We have found that nucGEMs are ejected from the nucleus during mitosis and very few nucGEMs remain upon nuclear reassembly. Subsequently, a slow accumulation of nucGEMs in daughter nuclei occurs, while nucGEMs presence in the cytosol slowly decreases^41^. In the example movie, we image nucGEMs in a cell where the particles are present in both the cytosol and nucleus, enabling us to quantify and compare their motion in both compartments. To separately analyze these compartments, we delineate the nucleus with the vital stain SiR-DNA (Figure 1b).

### Localization

To enable researchers to perform particle localization, we have provided a GUI interface to the *locate* function of the python package, Trackpy (**Locate** panel, Figure 2b). This function implements a classical algorithm developed by John Crocker and David Grier^33^. The input to the algorithm is a raw image with bright spots representing particles (Figure 2a). Dark spots can also be localized with image inversion: check the **Invert** checkbox in the panel (Figure 2b). We have localized GEM particles in the nucleus of the example movie (Figure 2c, single frame) over all frames and provided output that can be used for evaluation of results (Figure 2d,e). The first step is to preprocess the image by applying a dual bandpass filter (Figure 2a). A boxcar (rolling) average of specified size is subtracted from the image, which should be larger than the diameter expected for detecting spots but smaller than the distance between spots. This background subtraction is designed to remove long wavelength noise associated with variations in sensitivity of the camera’s pixels and uneven illumination^33^. The size for the boxcar filter (in pixels) is set with the parameter **Smoothing size** in the GEMspa localization panel (Figure 2b), and by default is set to one pixel more than the expected spot diameter. Next, the algorithm convolves the image with a gaussian kernel to remove high frequency electronic noise that can be introduced by the detector^33^. This is set with the **Noise size** parameter and by default is 1 px wide. Finally, a threshold is applied to the pre-processed image to clip all pixel values below a specified intensity value (the default is 1 but can be set with the **Threshold** parameter). If an image has been pre-processed with other tools, uncheck the **Preprocess** check box to skip all pre-processing. Next, the intensity maximas are found as the intersection of the image with its dilation, a morphological operation that sets a pixel value to the maximum within a given region. The **Separation** parameter is used here and corresponds to the minimum separation between particles. Maxima near the edge or too close to another maxima (within the separation distance) are dropped. Additionally, a percentile threshold is applied to eliminate “spurious” features by removing maximas with low brightness (**Percentile** parameter, default 64). The final step is to perform subpixel localization for each detected maxima. This is done with an iterative intensity-weighted centroid finding algorithm to refine the initial position estimate. The input to this algorithm is the typical diameter of a spot (**Diameter**), and the result can be evaluated by the subpixel bias metric (Figure 2d, panels 1&2). If the input diameter is set correctly, the decimal portions of the position estimates will be evenly distributed. However, if the diameter is too small, these histograms tend to be biased toward the edges (0 and 1)^33^. The uncertainty in the refined position estimations can be calculated from the intensity signal, the spot size (radius of gyration) and the expected diameter^50^ and is output by Trackpy (Figure 2d, final panel).

**Figure 2.**
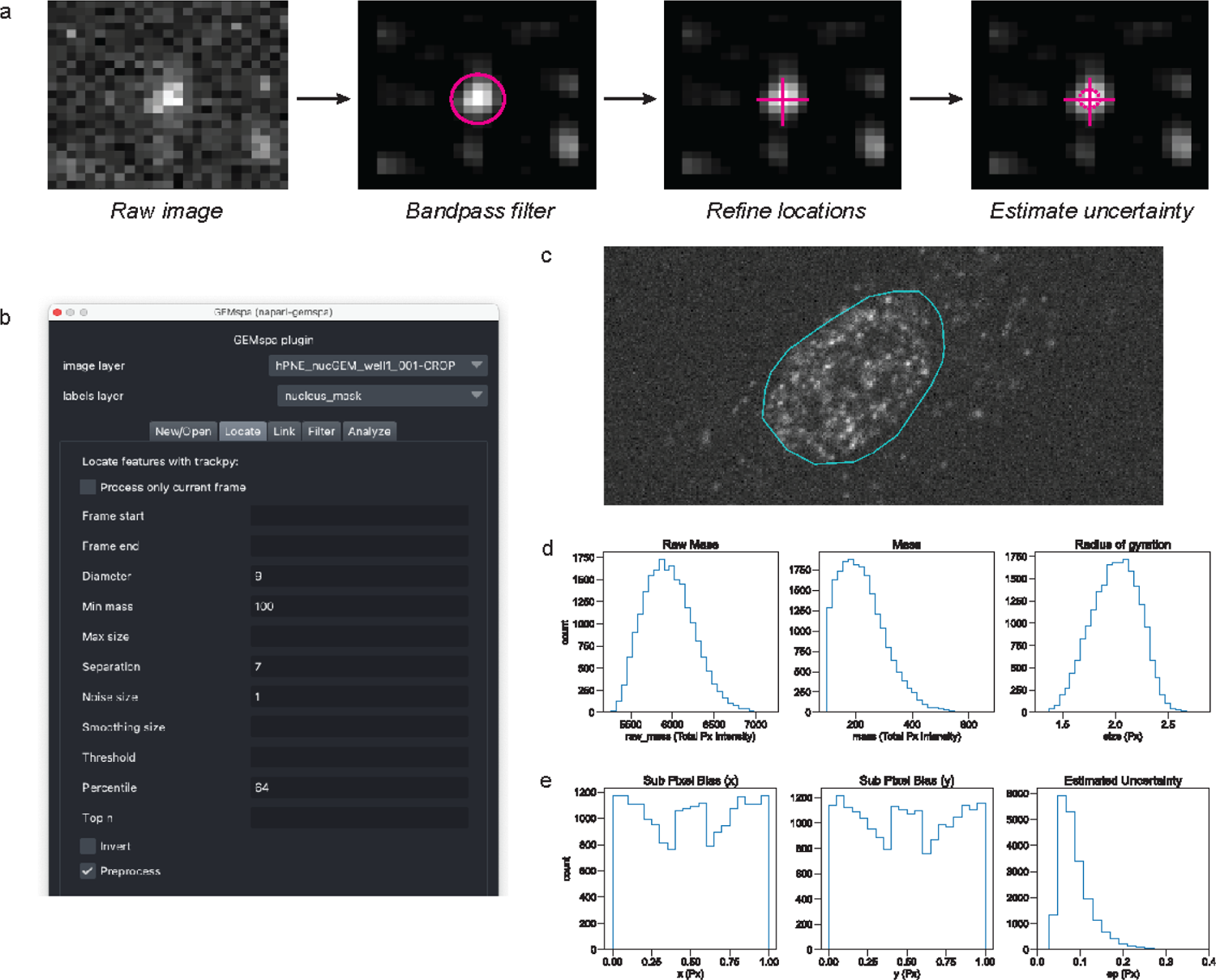
Overview of Localization with GEMspa. (a) Cartoons illustrating the steps of the particle localization algorithm. (b) A screenshot of the GEMspa plugin localization panel. (c) A screenshot of a single frame of the example GEMs movie showing nucGEM particles in an hPNE cell. (d) Histograms of Raw Mass, (Normalized) Mass, and Radius of Gyration output from the Trackpy particle localization algorithm for localized GEM particles within the cell nucleus (all frames). (e) Histograms of subpixel bias and estimated uncertainty output from Trackpy the particle localization algorithm for localized GEM particles within the cell nucleus (all frames).

This output can be used to aid in evaluation of the precision of subpixel localizations. The input parameters **Min Mass** (default 100) and **Max size** (no default) place additional constraints on what is considered a valid particle. Mass is defined as total integrated brightness and is highlighted by the <u>Trackpy documentation</u> as a “crucial parameter for eliminating spurious features.” The **Min Mass** input parameter is in normalized mass units, as converted from the raw mass of the input image when applying preprocessing filters. Both mass measurements are given in the output (Figure 2d). Size is the radius of gyration of the intensity (Figure 2d). Experimenting with these parameters can be useful for optimizing the localization algorithm to find valid features. GEMspa automatically displays subpixel bias and mass histograms to aid in parameter optimization. GEMspa outputs the results of localization as a new *points layer* in napari (Figure 2c), and includes the Trackpy output as properties of each point. To extract this data, save the layer as a tab/comma delimited text file with GEMspa (Supplementary files 2 & 3).

### Linking

To enable researchers to link particle positions into tracks, we have provided a GUI interface to the *link* function of the python package, Trackpy (**Link** panel, Figure 3a). This function implements the linking step from the Crocker and Grier algorithm^33^. The input to the algorithm is a table of particle positions for every frame of the movie (the output of the localization step). GEMspa currently provides access to only the linking algorithm that assumes Brownian motion, although Trackpy has expanded upon the original algorithm to allow for other types of motion. The linking algorithm assumes that the most likely position for a particle in the subsequent frame is its current position and attempts to find the combination of particle links that minimize the distance between the set of particles as a whole, allowing for new particles to appear and old particles to disappear at any time in the movie. As this can be computationally intensive due to the numerous possible combinations, the **Link range** parameter (Figure 3a) provides a maximum distance any particle can travel between frames of the movie. This should generally trim the possible combinations to a manageable number of subnets and allow for the algorithm to finish in a reasonable time. It is important to correctly set the link range large enough to link particles but small enough to keep the number of possible combinations under control. If particles are too dense and inter-particle spacing is less than the particle displacement in a single frame, tracking can become impossible. To effectively pick the link range in the example movie, we visualize both the distribution of particle displacements from one frame to the next as well as the distribution of inter particle spacing within each frame (Figure 3b). Prior to performing the linking step, the distances of particles in a frame to their nearest neighbors in the following frame can be taken as a proxy to determine the typical single particle displacement. Since two particles in one frame can move towards each other in the next frame, the typical displacement distance should be less than ½ of the interparticle spacing distance. The data from the GEM particles in the example movie show that the dual frame nearest neighbor estimate for single particle displacement is a bimodal distribution centered around 0.70px and 6.8px (medians of the two distributions split at x=4px) (Figure 3b). The larger distances likely represent new particles that appear throughout the movie. The ½ interparticle spacing ranges from 3.9-4.9px (25-75th percentile) with a minimum of 3.5px. Based on these results, the link range was set to 4px for linking. The **Memory** parameter allows particles to disappear for a set amount of frames and still be linked. Typically we do not allow missed frames when tracking GEMs, but Trackpy allows for this scenario. The **Min frames** parameter allows for filtering particles based on the number of frames tracked, which can be useful for only capturing tracks with sufficient length for proper statistics during the quantification step. After linking is performed, the distribution of post-linking particle displacements is no longer bimodal and matches the distribution that centers around 0.70px from the pre-linking approximation (Figure 3c). Another useful parameter to examine post-linking is the distribution of track lengths. This gives an idea of the length of time a typical GEM particle can be tracked. GEM particles can be lost due to photobleaching of the fluorophore and particles departing the focal plane.

**Figure 3.**
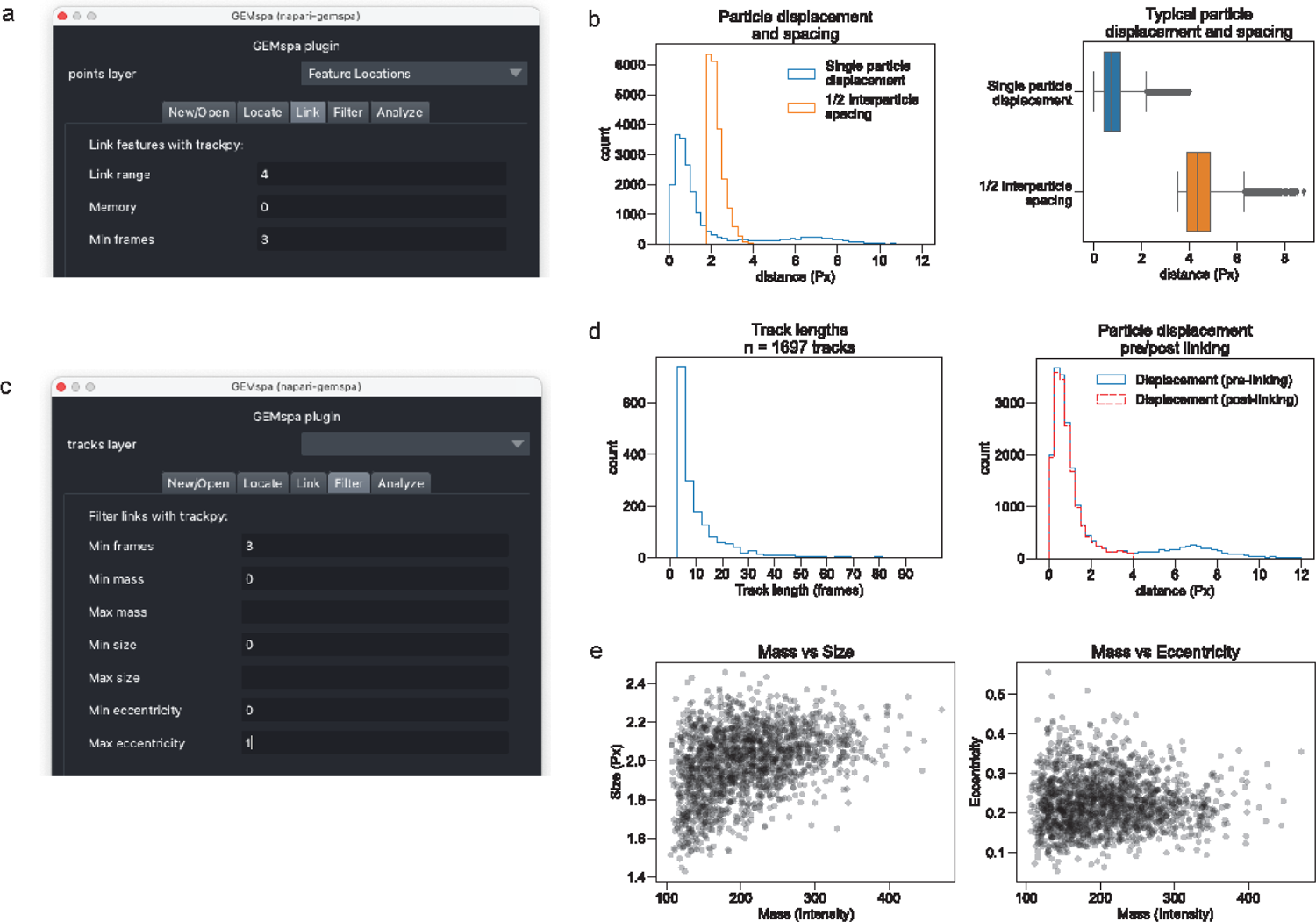
Overview of linking with GEMspa. (a) A screenshot of the GEMspa plugin linking panel (b) Histograms of estimated single particle displacements (pre-linking) and ½ interparticle spacing for localized GEM particles in the hPNE cell nucleus (left panel); Boxplots of same (right panel). (c) A screenshot of the GEMspa plugin filter links panel. (d) Histograms of linked GEM particles track lengths in the hPNE cell nucleus (left panel); Histograms of single particle displacements (pre and post-linking, right panel). (e) Scatter plots of average GEM particle mass vs. size (averaged over each track) (left panel) and average GEM particle mass vs. eccentricity (right panel).

In order to allow a filter step post-linking, GEMspa provides the **Filter** panel (Figure 3d). Once linking is complete, GEMspa displays two scatter plots with average particle properties over the length of the track (Figure 3e), and the track length histogram. Examining average particle properties over their lifetime provides more complete information on which to decide if the localized spot is a “true” particle. For example, particles with low mass may be out of focus and those with high eccentricity or size could be multiple particles or aggregates. Use the **Filter** panel to restrict entire tracks based on these properties (Figure 3d). For convenience, we also allow filtering of tracks by length at this stage.

GEMspa outputs the results of linking and filtering links as a new *tracks layer* in napari. The localization output is maintained in the properties layer and each localization is annotated with a particle ID. To extract the track data, save the layer as a tab/comma delimited text file with GEMspa (Supplementary files 4 & 5).

### Quantification

Once high quality particle traces have been obtained, the final step in analysis is to quantify the biophysical properties of the medium of interest. Here, we illustrate GEMspa’s quantification output (Figure 4a, **Analyze** panel) with our example movie and a comparison between GEMs in the nucleus and cytoplasm. We have used the *labels* layer of napari to draw masks for the nucleus (Figure 1b) and cytoplasm (not shown). GEMspa allows for selecting a labels layer for masking at either the particle localization step or the quantification step. We provide this function at the quantification step in case a researcher has imported a tracking file from outside software.

**Figure 4.**
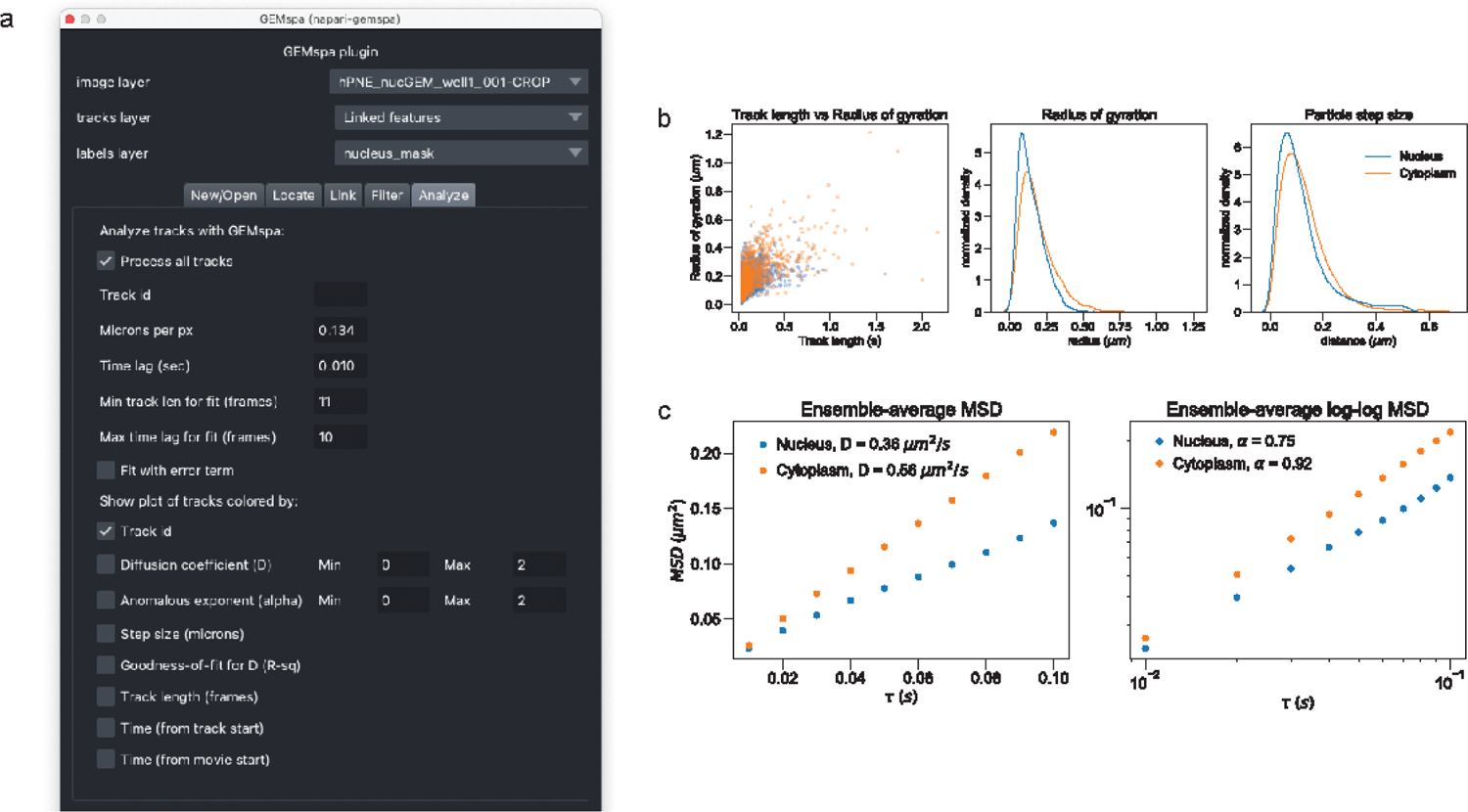
Quantification of average GEM particle motion with GEMspa. (a) A screenshot of the GEMspa plugin analysis panel. (b) Scatter plots of track length vs. radius of gyration for GEM particles in the nucleus and cytoplasm (left panel); normalized kernel density estimations of GEM particle radius of gyration distributions for particles in the nucleus and cytoplasm (middle panel); normalized kernel density estimations of GEM particle step size distributions for particles in the nucleus and cytoplasm (right panel). (c) Ensemble TA-(time average) MSD plots for GEM particles in the nucleus and cytoplasm illustration estimation of the effective diffusion coefficient (left panel); ensemble TA-MSD plots for GEM particles in the nucleus and cytoplasm on a log-log scale illustrating estimation of the anomalous exponent (right panel).

GEMspa provides a number of common output metrics generally used for quantification of SPT data. These metrics are saved as properties of each track in a new napari *tracks* layer upon running the analysis step. To extract the data, save the *tracks* layer with GEMspa (Supplementary Files 6 & 7). GEMspa also displays a table with summary information for each track (one row per track) as a pop-up window. Save this table as a text file to extract the data for further analysis.

The radius of gyration of a particle’s track quantifies the average distance a particle travels from its center of mass. This metric can help locate particles with little movement during their lifetime (Figure 4b). Outlier particles with long trajectories but very little movement may indicate stationary particles or aggregates. The step size (displacement) distribution is a good visualization for initial exploration of movement (Figure 4b), as it does not involve averaging and there are no assumptions regarding the type of motion. GEMs localized to the cytoplasm display a slight rightward shift to higher radius of gyration and step size values compared to nuclear GEMs at the 10 ms time scale. GEMspa calculates the (time-averaged) MSD for each trajectory and then averages over all particles at each time lag (Figure 4c), then fits this curve to the diffusion equation for Brownian motion to extract the effective diffusion coefficient. The data fit well to a linear increase in MSD with respect to time and we see on average faster diffusion of GEM particles in the cytoplasm compared to the nucleus (D = 0.36 µm^2^/s for nuclear GEMs in the nucleus and 0.56 µm^2^/s for cytoplasmic GEMs). To quantify anomalous motion, the exponent of a power law fit to the MSD can be extracted. In this case GEMspa extracts the exponent as the slope on the log-log scale for more consistent results (Figure 4c, second panel). GEM particles in the nucleus of this hPNE cell display increased subdiffusive motion (α = 0. 75) compared to cytoplasmic GEMs (α = 0. 92). The overall faster diffusion and less subdiffusive motion of GEMs in the cytoplasm compared to GEMs in the nucleus of this cell is consistent with bulk results we have previously reported^41^. This phenomena is opposed to what has been found for particles at a smaller size scale^51^ and could be explained by increased crowding and/or confinement specifically at the (40 nm) mesoscale in the nucleus compared to the cytoplasm.

GEMspa provides effective diffusion coefficient and anomalous exponent data on an individual track basis. It calculates the effective diffusion coefficient and anomalous exponent for each track and provides this data as tabular output as well as in the properties of a new tracks layer. The napari layers functionality can be used to color tracks by any quantity in the tracks layer properties (Figure 5a,b). The overlay of the tracks layer on another image (here, SiR-DNA) can help to visualize both the heterogeneity in particle motion and regions of exclusion. The previously reported exclusion of nucGEMs from heterochromatin and the nucleolus can be visualized in the overlay (Figure 5a).

**Figure 5.**
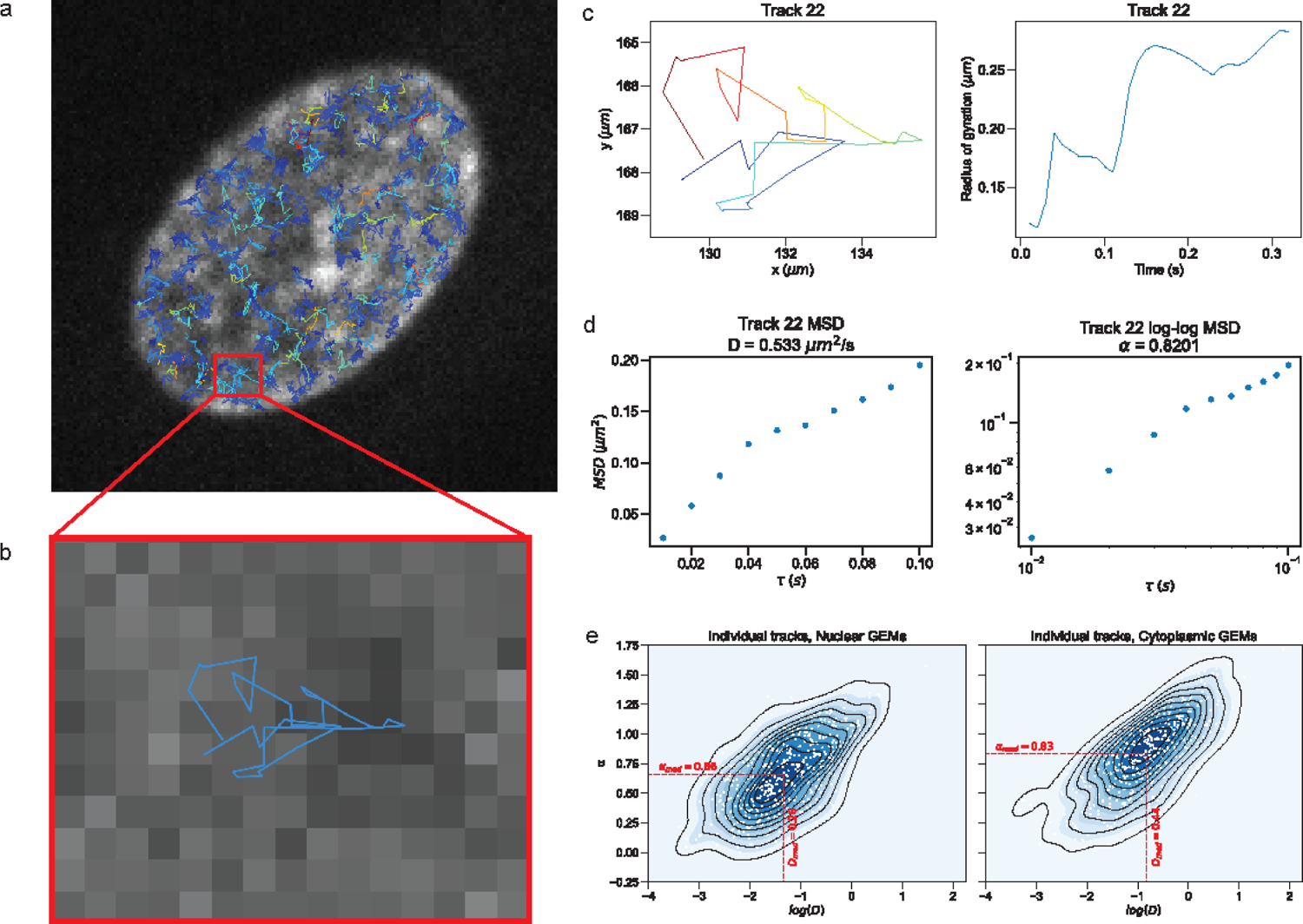
Quantification of individual GEM particle motion with GEMspa. (a) Screenshot of napari tracks layer visualizing individual tracks colored by effective diffusion coefficient. (b) Zoom-in view of an individual track (ID=22). (c) Plot of an individual track (ID=22) colored by time (left panel); Plot of the radius of gyration over time for track 22 (right panel). (d) TA-(time average) MSD plot for track 22 illustrating estimation of the effective diffusion coefficient (left panel); TA-MSD plot for track 22 on a log-log scale illustrating estimation of the anomalous exponent (right panel). (e) Bivariate kernel density estimates with overlaid scatter plots of GEM particles (log scale) effective diffusion coefficient and anomalous exponent for GEMs in the nucleus (left panel) and cytoplasm (right panel).

GEMspa provides an option to visualize the data for an individual track in depth. Uncheck **Process all tracks** (Figure 4a) and enter a track ID to view the path of the selected track (Figure 5c). The full path of the track is displayed in addition to the radius of gyration at each time step (Figure 5c). Examining this radius of gyration plot can help visualize the nature of a track’s motion as it evolves over time. GEMspa also displays the MSD data for the track. This can be helpful to understand the type of motion that gives rise to the shape of the MSD curve. In the example track, the slope of the MSD changes at larger time-lags and fits less well to the diffusion equation with linear scaling than it does with log-log scaling (r^2^ *of the linear fit = 0.87*, r^2^ *of the log* − *log fit* = 0. 97). The effective diffusion coefficient can be extracted, but care should be taken to quantify how well the model for Brownian motion agrees with the track data. In this case, the track’s motion is more consistent with a subdiffusive model (α = 0. 82) (Figure 5d).

The summary output of GEMspa for each track can be used to understand the heterogeneity of GEM motion in the cell. The heatmaps (Figure 5e) show the bivariate distribution of effective diffusion coefficients and anomalous exponents for each particle in the cytoplasm and nucleus. We can see an overall trend of increased diffusive motion in the cytoplasm compared to the nucleus (median D = 0.26 µm^2^/s for nuclear GEMs in the nucleus and 0.44 µm^2^/s for cytoplasmic GEMs) and more subdiffusive motion in the nucleus (median α = 0. 66 for nuclear GEMs and median α = 0. 83 for cytoplasmic GEMs), consistent with the ensemble results (Figure 4c), however we can also see a high level of diversity in particle motion in both compartments. The individual track variability of the effective diffusion coefficient is lower for nuclear GEMs (std-dev of D for nuclear GEMs = 0.34 and cytoplasmic GEMs = 0.52 µm^2^/s) while the variability for the anomalous exponent is similar in both compartments (std-dev of α for nuclear GEMs = 0.31 and cytoplasmic GEMs = 0.32). Here we report trends but do not attempt to calculate statistical tests based on data from a single cell. However, these results are consistent with measurements we have previously reported for GEM particles at the mesoscale in the 100 ms time scale^41^.

If the checkbox **Fit with error term** (Figure 4a) is selected, GEMspa will extract the effective diffusion coefficient from a diffusion equation that includes an y-intercept term. The MSD data will cross the y-axis at the x (time-lag)=0 point above y (MSD)=0. This can occur since localization error is inherent in the methods used to pinpoint the particle position with sub-diffraction accuracy. Care should be taken to examine the distribution of diffusion coefficients in this case as it can result in negative values when the data does not fit well to a linear model. These types of tracks generally correspond to tracks with low goodness-of-fit (r^2^) values when fitting the equation without the y-intercept term (data not shown).

Finally, GEMspa allows the researcher to choose to output all tracks (over all time frames) on a single plot. This is similar to coloring the tracks layer in napari by properties related to track motion, but provides an alternate format for saving as a figure and a few extra options for display (Figure 6).

**Figure 6.**
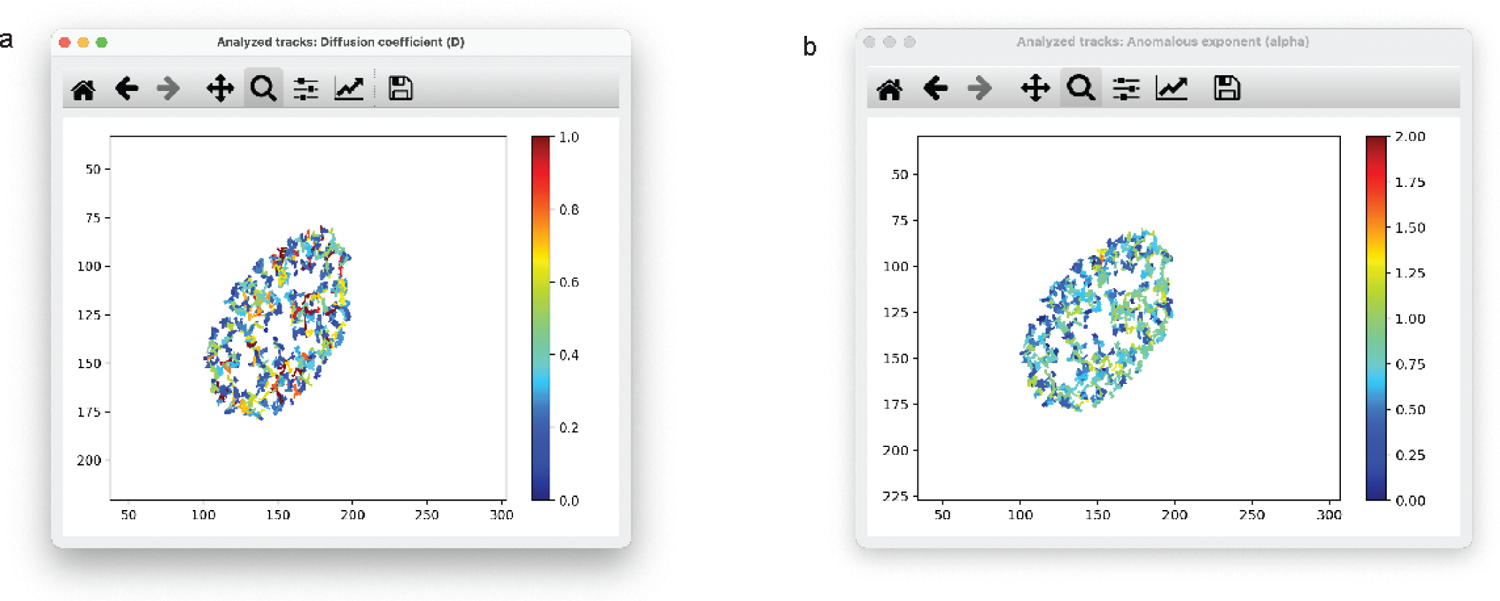
GEM particle tracks colored by parameters quantifying motion. (a) GEM particle tracks in the nucleus of the hPNE cell colored by effective diffusion coefficient. (b) GEM particle tracks in the nucleus of the hPNE cell colored by anomalous exponent.

## DISCUSSION

Here we present GEMspa, a napari plugin that allows a researcher to perform analysis of data from single particle tracking microrheology experiments with an easy to use GUI interface for all aspects of the analysis workflow. It is presented as a series of tab-separated plugin modules to perform localization, tracking, filtering of tracks and quantification of motion. The design goal for GEMspa was to unify analysis into a single application and provide visual feedback to enable the user to optimize parameter settings at each step of the workflow. A number of algorithms exist for particle localization and tracking and GEMspa does not directly implement a new algorithm for these steps. Instead it provides an interface to a classical algorithm that is well established and can be applied when knowledge of the expected type of motion is limited. In the future, we aim to implement interfaces for additional localization/tracking algorithms to allow the researcher to choose the implementation that provides the best results for their data. Performing exploratory analysis for individual SPT movies is a necessary first step to find the optimal settings needed for batch processing. The interactive nature of the GEMspa GUI combined with the napari tracks layer visualization capabilities will allow the researcher to understand the nature of their data and optimize parameter settings both at each step individually and as a whole. Instead of performing localization and tracking separately and then implementing quantification afterwards, the quantification step can be integrated into the analysis workflow in real time. The suite of quantification functions provide instant results for those that may not have the computation skills to easily extract more advanced results from motion models. There are many methods that have been applied to quantify SPT tracking data, and GEMspa provides output from models that are the most common and straightforward to interpret. In the future, more quantification methods can be added. For example, the ergodicity breaking parameter^38, 41^ is a common method to check if there is intrinsic variability in the motion of individual particles in an ensemble. This can be caused by non-specific interactions with or heterogeneity in the biophysical properties of the cellular environment. Another useful measure to probe for anomalous motion is to examine the distribution of directional change for particle trajectories at successive time intervals^52^. In a pure random walk, these angles would be distributed evenly between 0 degrees (continuing in the same direction) and 180 degrees (reversing direction), while a skewed distribution could indicate confined or directed motion.

Some limitations exist when analyzing SPT data. First, particle localization and tracking are subject to error. The position of a particle can only be determined to a certain level of precision. This error can lead to underestimation of the anomalous exponent^52, 53^. Tracking errors can occur especially when particles are dense and the algorithm incorrectly connects two different particles in successive frames. Second, the statistical noise due to the inherent variability of random motion needs to be overcome. Usefulness of the MSD depends on having enough data over which to average in order to reduce the variance and quantify motion with rigorous statistics. In the case of the time-averaged MSD, this means longer tracks are better, since this increases the number of time increments over which to take the average. Care should be taken to examine the distribution of track lengths and decide how to limit the number of time-lags over which to fit the MSD to extract the diffusion coefficient and anomalous exponent. Removal of very short tracks that have little data for averaging is also advisable.

The GEMspa napari plugin is designed for initial data exploration, parameter optimization and visualization. While it can be used for analysis of single SPT movies, processing many movies (a “batch” mode) is not yet possible. Future work will be centered around allowing the export of parameters and settings and an additional GUI to process entire data sets and output composite results that can be used for rigorous statistical analysis. We have an <u>existing implementation</u> for batch processing that is actively used in our lab and is being rolled over to the napari plugin format.

In conclusion, GEMspa is a comprehensive, user-friendly napari plugin designed to streamline the analysis of single particle tracking microrheology data, providing real-time quantification and interactive visualization.

## METHODS

### Experimental Methods

Detailed methods for the hPNE cell line, plasmid construction, lentivirus production and cell transduction have been previously described^7,41^. For the SPT movie, nucGEMs were imaged at 100 Hz with a spinning disk confocal as previously reported ^41^.

### Analysis of GEMs in single hPNE cell

Segmentation of the nucleus was based on the SiR-DNA stain; for the cytoplasm based on a max-projection of the GEMs movie, subtracting the nuclear region. The GEMspa plugin was used to perform particle localization and tracking (with Trackpy), and the analysis of motion. The GEMspa plugin (<u>napari-gemspa</u>) can be installed from PyPi with pip or through the napari Plugins menu.

### Details of plugin implementation

The plugin is implemented in python. It basically provides a GUI interface for localization and tracking to the respective Trackpy methods. The only additional processing is to filter localized particles based on a mask from the labels layer of napari. The functions for quantification of motion are implemented in a python package developed in-house and available on PyPi (<u>gemspa-spt</u>). Filtering of track data at the analysis step with a mask results in the removal of all tracks that contain points that are outside labeled regions. The GEMspa plugin home page provides a <u>detailed user guide</u> with usage instructions and a description of all analysis methods.

## Acknowledgements

We would like to thank Daniel Elnatan for his input on plugin design and sharing of code, and Gururaj Rao Kidiyoor for sharing the example SPT movie that was used to illustrate the plugin.

## Notes

### Competing Interest Statement

The authors have declared no competing interest.

