## Supplementary figures and images for "GEMspa: a Napari plugin for analysis of single particle tracking data"

### SupplementaryMovie

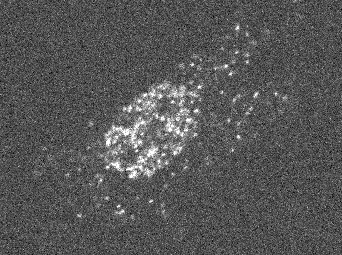
